# Impaired peri-olfactory cerebrospinal fluid clearance mediated cognitive decline after dyssomnia

**DOI:** 10.1101/2022.03.05.482807

**Authors:** Ying Zhou, Wang Ran, Zhongyu Luo, Jianan Wang, Mengmeng Fang, Kai Wei, Jianzhong Sun, Min Lou

## Abstract

Animal experiments have demonstrated the dependency of CSF clearance function on age and sleep, which partially underlay the cognitive decline in the elderly. However, evidence is lacking in humans mainly due to limited method to assess CSF clearance function. We aimed to image CSF clearance pathways in human brain by dynamic MRI with intrathecal contrast agent as a CSF tracer. We performed T1-weighted and T2-fluid attenuated inversion recovery imaging with equal sequence parameters before and at multiple time points after intrathecal injection of contrast agent to visualize the putative meningeal lymphatic pathway, peri-olfactory nerve pathway and peri-optic nerve pathway. We defined the CSF clearance function as the percentage change in signal unit ratio of critical locations in these pathways from baseline to 39 hours after intrathecal injection. CSF clearance through the putative meningeal lymphatic and peri-neural pathways were clearly visualized in all 85 patients. The CSF clearance function of meningeal lymphatic and peri-neural pathways were reduced with aging (all *P* < 0.05). The CSF clearance function through peri-olfactory nerve pathway was positive correlated with sleep quality and cognitive function (both *P* < 0.05). Moreover, the CSF clearance function through peri-olfactory nerve pathway mediated the association of sleep quality with cognitive function (percent change in *β* [bootstrap 95% confidence interval]: 35% [−0.220, −0.003]). The study shows promise for dynamic MRI with intrathecal injection of contrast agent as a method to assess CSF clearance function through putative peri-neural pathways, and interprets the impaired clearance through putative peri-olfactory nerve pathway may explain the cognitive decline in patients with dyssomnia.

## Introduction

CSF is considered to be produced primarily by the choroid plexuses and circulates through the subarachnoid space and ventricles to maintain a stable environment for the brain and spinal cord.^1^ Traditionally, it was accepted that CSF mainly drain through arachnoid granulations from the subarachnoid space to the dural venous sinuses.^2^ However, this concept has been increasingly challenged in recent years. Accumulating evidences from various species including rodent, artiodactyla and nonhuman primate have shown that tracers injected into the CSF can be found within lymphatic vessels from the cranium and spine, as well as along exiting cranial and spinal nerves.^3,4^ In addition, it is worth noting that several animal studies have found the dependency of CSF clearance function on age and sleep, which may partially underlie the cognitive decline in the elderly.^5–7^

Recently, our group and another research group have independently visualized the efflux of CSF through parasagittal dura, reflecting the putative meningeal lymphatic pathway in humans.^8,9^ However, the evidences of peri-neural pathways are relatively limited and controversial. A PET study using radionuclide tracers reported nasal turbinate as a location of the CSF drainage pathway in healthy human and Alzheimer’s disease patients.^10^ Conversely, a MRI study found that gadolinium injecting into CSF as tracer could not be detected on nasal turbinate in patients with normal pressure hydrocephalus.^11^ More recently, based on five participants, another MRI study suggested that all peri-neural spaces surrounding the cranial nerves were filled with CSF.^12^ Thus, since the existence of peri-neural pathways in humans is still controversial, rare study reported the function of these pathways, as well as their potential interactions with sleep quality and cognitive function.

In the current study, by utilizing dynamic brain 3-dimensional T1-weighted imaging and high-resolution 2-dimensional T2-fluid attenuated inversion recovery imaging with intrathecal injection of a gadolinium contrast agent, we visualized three main CSF draining pathways including (1) putative meningeal lymphatic pathway through parasagittal dura; (2) putative peri-olfactory nerve pathway through subarachnoid space adjacent to straight gyri and superior turbinate, middle turbinate and inferior turbinate; and (3) putative peri-optic nerve pathway through middle intraorbital segment of the optic nerve and corneoscleral part. We then assessed the CSF clearance function in each pathway and further investigated their relationships with age, sleep quality and cognitive function.

## Materials and methods

### Experimental Design and Study Subjects

In this prospective observational study, we enrolled consecutive patients with indications for lumbar puncture and voluntary participation. Exclusion criteria were known adverse reaction to contrast agents, history of severe allergic reactions in general, renal dysfunction, and pregnant or breastfeeding females. All patients underwent MRI before and at multiple time points including 4.5 hours, 15 hours, and 39 hours after intrathecal injection of gadodiamide during a study period from April 2018 to October 2021. The Telephone Montreal Cognitive Assessment (T-MoCA) was used to evaluate the cognitive function from five domains including attention and calculation, language, abstraction, delayed recall, and orientation over the phone in November 2021. The total score was 22 points with higher scores indicating better cognitive function.^13^ The Pittsburgh Sleep Quality Index (PSQI) was used to evaluate the quality and patterns of sleep from seven domains including subjective sleep quality, sleep latency, sleep duration, habitual sleep efficiency, sleep disturbances, use of sleeping medication, and daytime dysfunction during hospitalization (at the same time as MRI). The total score range from 0 to 21 with higher scores indicating more severe dyssomnia.^14^ The study, including the administration of intrathecal gadolinium agents, obtained approval from the local human ethics committee. All clinical investigation was conducted in compliance with the principles expressed in the Declaration of Helsinki. All patients were included after informed consent.

### MRI protocol

A 3.0T MRI scanner (GE 750; GE Healthcare, Chicago, IL) with equal imaging protocol settings at all time points was applied to acquire axial head 3-dimensional T1-weighted, high-resolution coronal head 2-dimensional T2-fluid attenuated inversion recovery and axial neck T1 fat-suppression imaging. The main imaging parameters were, for axial head 3-dimensional T1-weighted: repetition time = 7.3 milliseconds, echo time = 3.0 milliseconds, flip angle = 8, thickness = 1 mm, field of view = 25 × 25 cm^2^, matrix = 250 × 250 pixels; for high-resolution coronal head 2-dimensional T2-fluid attenuated inversion recovery: repetition time = 8,400 milliseconds, echo time = 152 milliseconds, flip angle = 90°, thickness = 3 mm, no slice gap, field of view = 18 × 18cm^2^, matrix = 320 × 320 pixels; and for neck T1 fat-suppression imaging: repetition time = 535 milliseconds, echo time = 9.1 milliseconds, thickness = 4 mm, slice gap = 0.5 mm, field of view = 22 × 22cm^2^, matrix = 130 ×100 pixels.

### Intrathecal Administration of Gadodiamide

The site of intrathecal injection of the contrast agent is L3-4 or L4-5 lumbar intervertebral space. Intrathecal injection of 1 ml of 0.5 mmol/ml gadodiamide (Omniscan; GE Healthcare) was preceded by verifying the correct position of the syringe tip in subarachnoid space in terms of CSF backflow from the puncture needle^8^. Following needle removal, patients were instructed to rotate themselves around the long axis of the body twice, and then remain in the supine position until 4 hours after intrathecal injection.

### Evaluation of Imaging

A neuroradiologist (R.W.) with 5 years of experience and blind to clinical and other image data independently placed regions of interest at all time points using RadiAnt (Medixant, Poznan, Poland), a Digital Imaging and Communications in Medicine viewer. We placed regions of interest at middle intraorbital segment of the optic nerve (hereinafter referred to as the “optic nerve”) and corneoscleral part on axial head 3-dimensional T1-weighted, parasagittal dura, subarachnoid space adjacent to straight gyri (hereinafter referred to as “adjacent straight gyri”), superior turbinate, middle turbinate and inferior turbinate on high-resolution coronal head 2-dimensional T2-fluid attenuated inversion recovery, and deep cervical lymph node on neck T1 fat-suppression imaging.

We measured mean signal unit for each region of interest and normalized them against references. According to previous studies, we chose vitreous body of the ocular bulb on axial head 3-dimensional T1-weighted imaging and high-resolution coronal head 2-dimensional T2-fluid attenuated inversion recovery imaging, and medial pterygoid muscle on neck T1 fat-suppression imaging as references, since there was no significant tracer accumulation in these regions after intrathecal injection of gadodiamide.^8,11,15^ For each time point, we determined the signal unit ratio between the regions of interest and references. We defined the clearance function of each region as the percentage changes in signal unit ratio from baseline to 39 hours, with lower percentage changes indicating better clearance function.

In each patient, we compared the signal unit ratio at 4.5 hours, 15 hours, and 39 hours at each region, and the time point of relative high signal unit ratio is defined as the peak time point of CSF tracer enrichment. We then compared the peak time point among different regions, and illustrated them as the tracer appeared at one region before/at the same time point as/later than another region.

### Statistical Analysis

Continuous data were described as mean with standard deviation or median with interquartile spacing. Categorical data were presented as numbers and percentage. Differences between continuous or categorical data in the same individual were determined using paired-sample *t* tests or paired-sample Wilcoxon signed rank test. Correlations between continuous or rank data were determined using Pearson or Spearman correlation analysis. Multivariable analyses were determined using linear regression analysis. Differences between continuous data were determined using linear mixed models with a random intercept.

Mediation analyses were used to assess whether the association between sleep (PSQI total scores) and cognitive function (T-MoCA total scores) could be mediated by the CSF clearance function. To perform mediation analysis,^16^ we tested 3 pathways: step 1, the association of sleep with cognitive function; step 2, the association of sleep with the CSF clearance function; and step 3, the association of the CSF clearance function with cognitive function, controlling for sleep. If all 3 associations are satisfied, the indirect effect established. The extent of indirect effect is estimated by examining the change in the *β* of PSQI total scores for T-MoCA total scores before and after including the CSF clearance function. We also adopted the product of coefficient approach for mediation analysis. We calculated the 95% confidence interval of the indirect effect using bootstrapping technique (5,000 replicate samples with replacement). Statistical analyses were performed using the SPSS software version 22.0 (IBM, Armonk, NY). Statistical significance was accepted at the 0.05 level (2-tailed).

## Results

### Patients

A total of 85 patients (43 males, mean age = 59 years, age range = 21-85 years; 42 females, mean age = 56 years, age range = 18-79 years) were enrolled. The diagnosis included peripheral neuropathy (*n* = 43), neurodegenerative diseases (*n* = 17), encephalitis (*n* = 16), normal pressure hydrocephalus (*n* = 4), suspected CSF leakage (*n* = 4), and hepatic encephalopathy (*n* = 1). T-MoCA scores averaged 14.2 ± 4.2 and PSQI scores averaged 6.7 ± 4.2. Supplementary Table 1 shows the basic clinical data.

### Tracer enrichment in all predefined pathways

In all patients, the signal unit ratio of parasagittal dura, adjacent straight gyrus, superior turbinate, middle turbinate, inferior turbinate, optic nerve and corneoscleral part changed significantly after the gadolinium injection (Fig. 1; Fig. 2). We present the peak time point at each region among all patients (Supplementary Table 2). In summary, the CSF tracer appeared at adjacent straight gyrus before or at the same time point as the superior, middle and inferior turbinate in 86.3% (44/51) of patients, whereas later than either of the superior, middle and inferior turbinate in 13.7% (7/51) of patients. In addition, the tracer appeared at the optic nerve before or at the same time point as corneoscleral part in 90.2% (74/82) of patients, whereas later than corneoscleral part in 9.8% (8/82) of patients. Additionally, the tracer appeared at the deep cervical lymph node later than or at the same time point as the parasagittal dura, adjacent straight gyrus, superior turbinate, middle turbinate, inferior turbinate, optic nerve and corneoscleral part in 80% (12/15) of patients, whereas early than adjacent straight gyrus and superior turbinate in one patient, optic nerve and corneoscleral part in one patient, and parasagittal dura and corneoscleral part in one patient. Supplementary Table 3 shows the proportion of different peak time points at each region among all patients. The peak time point in the superior, middle and inferior turbinate were significantly later than adjacent straight gyrus (all *P* < 0.05), but there was no significant difference of peak time point between optic nerve and corneoscleral part (*P* = 0.237).

**Figure 1.**
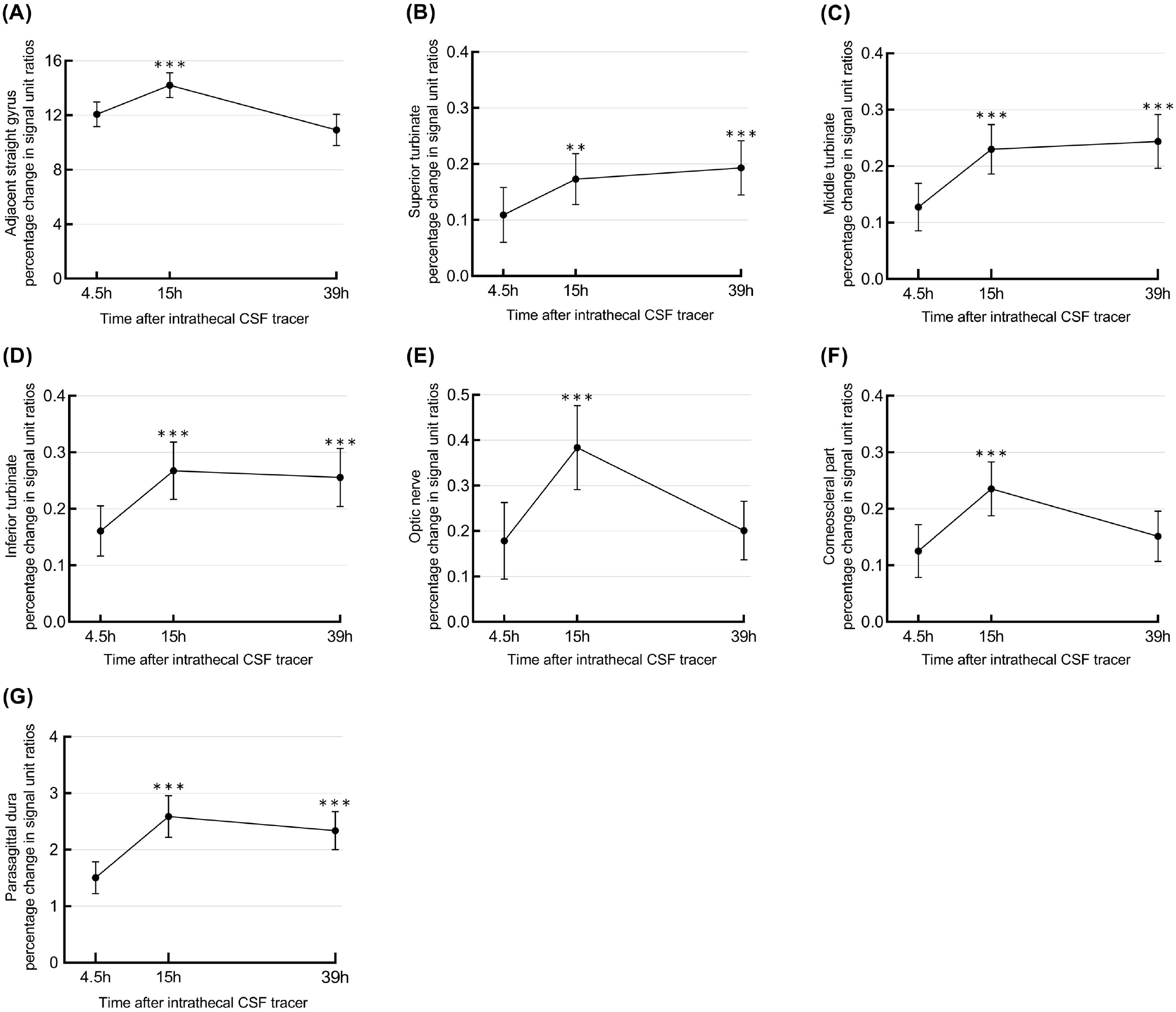
Tracer enrichment in different locations. The percentage change in signal unit ratios over time from CSF tracer administration through adjacent straight gyrus (A), superior turbinate (B), middle turbinate (C), inferior turbinate (D), optic nerve (E), corneoscleral part (F), and parasagittal dura (G). Error bars refer to 95% confidence interval. The signal change was highly significant in all locations (linear mixed model analysis; **P < 0.01, ***P < 0.001).

**Figure 2.**
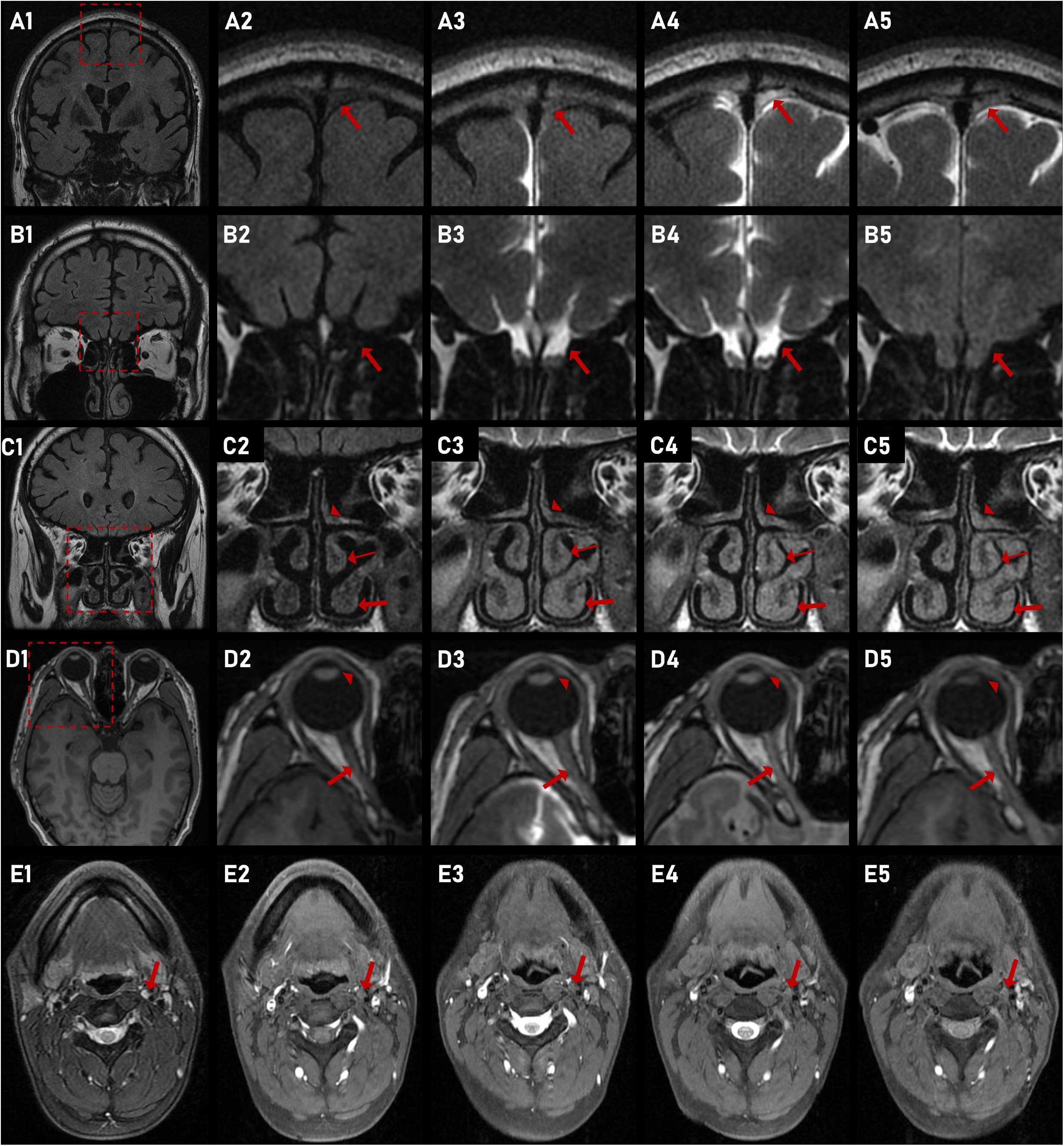
Representative images of putative meningeal lymphatic and peri-neural pathways and deep cervical lymph nodes. The clearance of putative meningeal lymphatic pathway and peri-olfactory nerve pathway on images of head high-resolution 2-dimensional T2-fluid attenuated inversion recovery imaging (A, thick arrow refers to parasagittal dura, B, thick arrow refers to adjacent straight gyrus, C, triangle, thin and thick arrow refer to superior, middle and inferior turbinate, respectively), putative peri-optic neural pathway on head 3-dimensional T1-weighted imaging (D, triangle and thick arrow refer to corneoscleral part and optic nerve) and deep cervical lymph node on neck T1 fat-suppression imaging (E, thick arrow refers to deep cervical lymph node) are shown, respectively. A1-E1, A2-E2, A3-E3, A4-E4 and A5-E5 were baseline images, corresponding magnified images within the red box on A1-E5, 4.5 hours images, 15 hours images and 39 hours images after the intrathecal administration of gadodiamide, respectively.

### Impairment CSF clearance function with aging

The worse clearance function of parasagittal dura, adjacent straight gyrus, superior turbinate and middle turbinate were significantly related to aging (all *P* < 0.05; Fig. 3). However, no significant correlations were found between the clearance function of inferior turbinate, optic nerve and corneoscleral part and aging (all *P* > 0.05; Fig. 3). In addition, when age and neurodegenerative disease were set as independent variables in linear regression, aging was still significantly associated with the worse clearance function of the parasagittal dura (*β* = 0.046, *P* = 0.001), adjacent straight gyrus (*β* = 0.153, *P* = 0.001), superior turbinate (*β* = 0.007, *P* = 0.001) and middle turbinate (*β* = 0.004, *P* = 0.036). The representative images in Fig. 4 and Fig. 5 show the relationship between the clearance function of parasagittal dura, superior turbinate, middle turbinate and inferior turbinate and aging.

**Figure 3.**
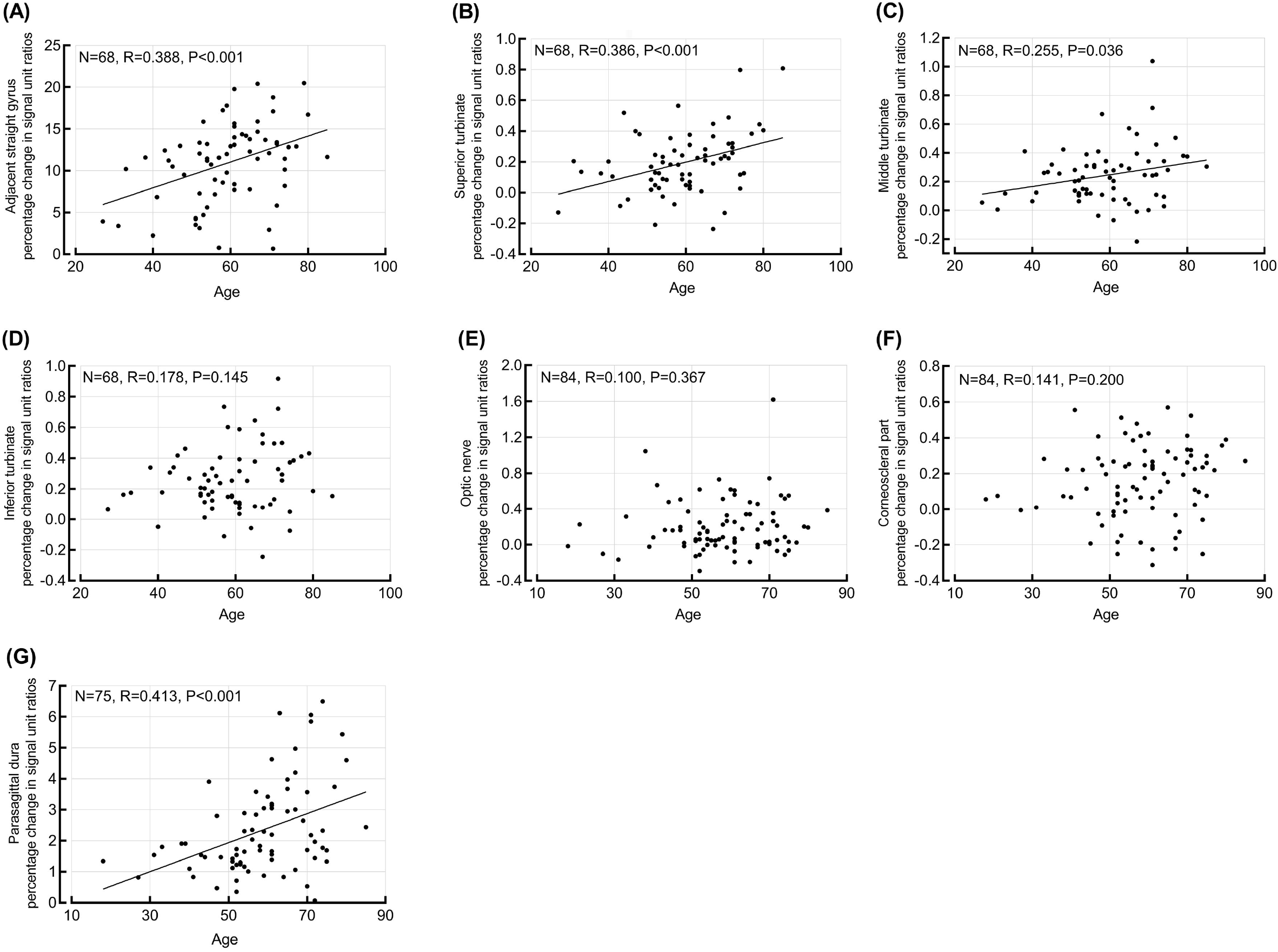
Correlations between age and the clearance function on the seven predefined locations. There was a positive correlation between age and the clearance function through adjacent straight gyrus (A), superior turbinate (B), middle turbinate (C), or parasagittal dura (G), respectively. No significant correlation was observed in the clearance of inferior turbinate (D), optic nerve (E), and corneoscleral part (F) with age. Each plot shows the sample size, fit line, and the Pearson correlation coefficient (R) with P-value.

**Figure 4.**
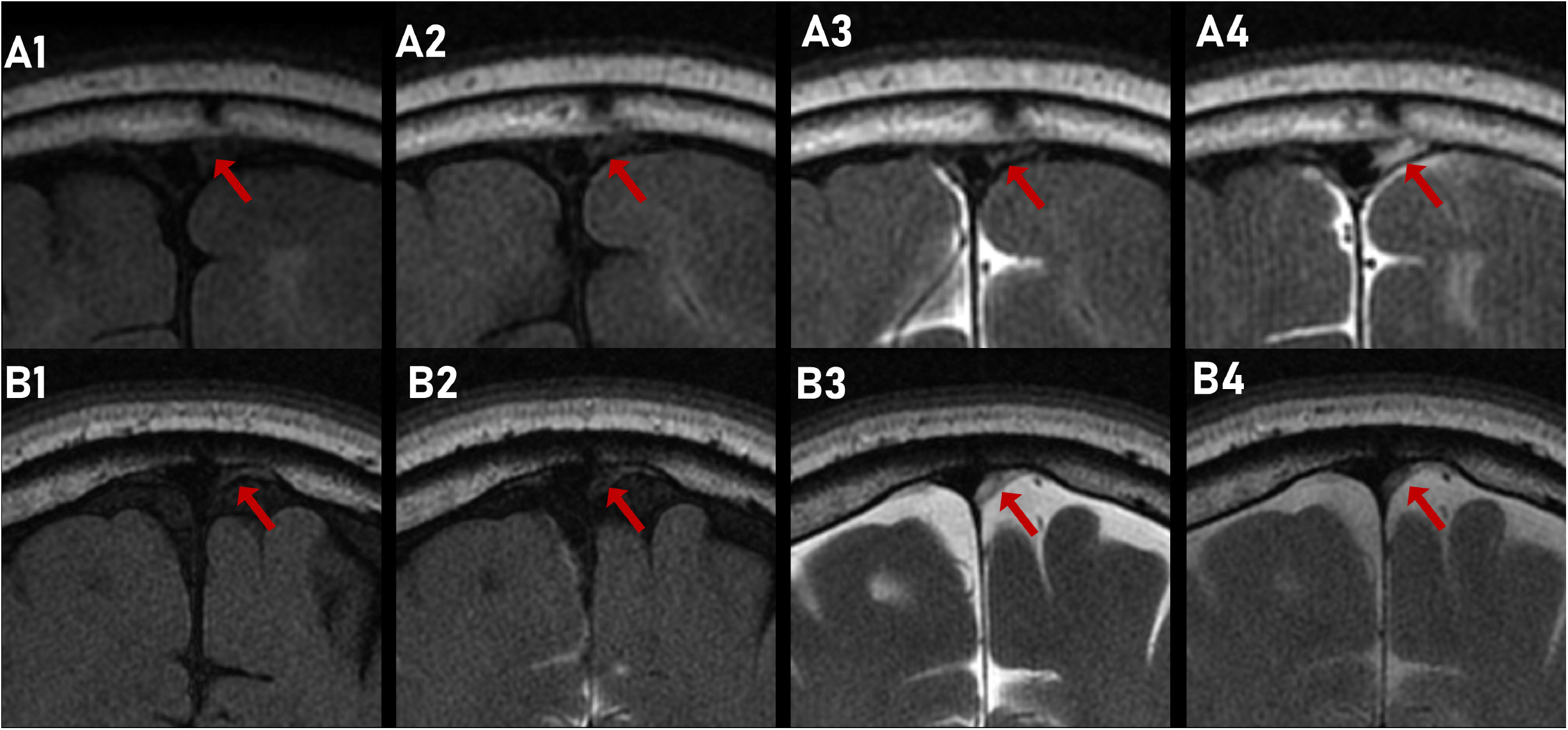
Representative images of the putative meningeal lymphatic pathway in two patients. From the images of coronal head 2-dimensional T2-fluid attenuated inversion recovery at baseline (A1 and B1), 4.5 hours (A2 and B2), 15 hours (A3 and B3), and 39 hours (A4 and B4) after the intrathecal administration of gadodiamide, the signals of parasagittal dura (arrow) changed significantly, indicating the CSF clearance through putative meningeal lymphatic pathway. The peak time point of signal unit ratio in the 65-year-old patient (at 39 hours, A4) is later than that in the 31-year-old patient (at 15 hours, B3).

**Figure 5.**
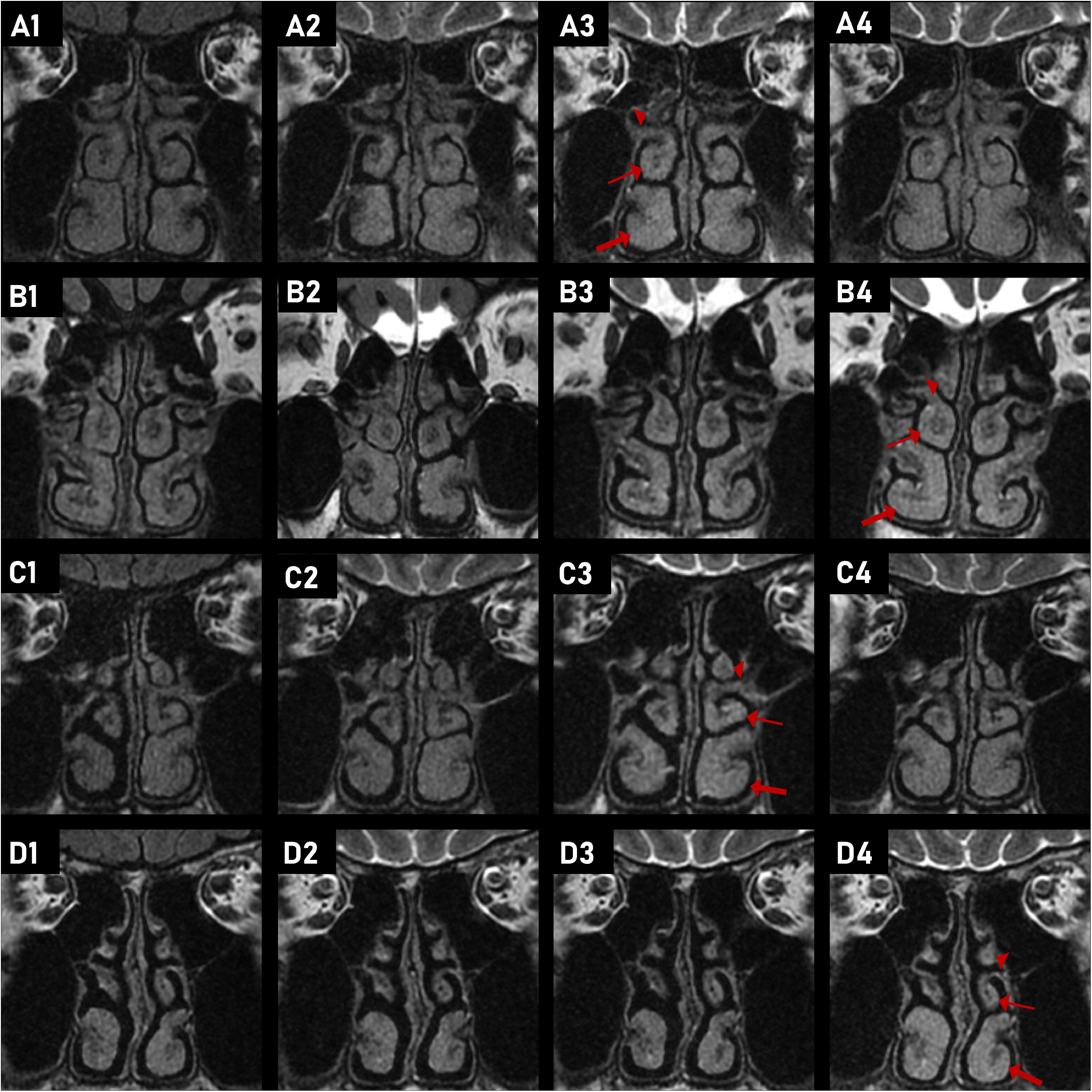
Representative images showing the delayed clearance of the putative peri-olfactory nerve pathway in aging patients and patients with poor sleep quality. The signals of the turbinates (triangle, thin and thick arrow refer to superior, middle and inferior turbinate, respectively) changed significantly from images of coronal head 2-dimensional T2-fluid attenuated inversion recovery at baseline (A1, B1, C1 and D1), 4.5 hours (A2, B2, C2 and D2), 15 hours (A3, B3, C3 and D3), and 39 hours (A4, B4, C4 and D4) after the intrathecal administration of gadodiamide, indicating the CSF clearance. On all turbinates, the peak time point of signal unit ratio in the 54-year-old patient (at 15 hours, A3) is earlier than that in the older 79-year-old patient (at 39 hours, B4). The peak time point of signal unit ratio in the 60-year-old patient with good quality of sleep (Pittsburgh Sleep Quality Index score of 1) (at 15 hours, C3) is earlier than that in the 63-year-old patient with poor quality of sleep (Pittsburgh Sleep Quality Index score of 6) (at 39 hours, D4).

### Relationship between CSF clearance and cognitive function

The clearance function of superior turbinate, middle turbinate and inferior turbinate were positively realted to T-MoCA total scores (all *P* < 0.05; Table 1). After adjusting for age and education years, the clearance function of superior turbinate, middle turbinate and inferior turbinate were independent protective factors of T-MoCA total scores (*β* = −5.359, *P* = 0.033; *β* = −7.206, *P* = 0.009; *β* = −5.997, *P* = 0.006). For the sub-item of T-MoCA, the clearance function of superior turbinate was positively correlated with delayed recall, the clearance function of middle turbinate was positively correlated with attention and calculation, language, abstraction and orientation, and the clearance function of inferior turbinate was positively correlated with attention and calculation and language (all *P* < 0.05; Table 1). However, no significant relationship was found between T-MoCA scores and the clearance function of parasagittal dura, adjacent straight gyrus, optic nerve and corneoscleral part (all *P* > 0.05; Table 1).

**Table 1.** Correlations between cognitive function and percentage change of signal unit ratios from baseline to 39 hours in predefined seven locations.

|  |  | T-MoCA<br>total<br>scores | Attention<br>and<br>calculation | Language | Abstraction | Delayed<br>recall | Orientation |
| --- | --- | --- | --- | --- | --- | --- | --- |
| Parasagittal dura<br>(n = 51) |  |  |  |  |  |  |  |
|  | R | -0.113 | -0.067 | -0.108 | -0.094 | -0.070 | -0.050 |
|  | P | 0.432 | 0.639 | 0.451 | 0.511 | 0.625 | 0.725 |
| Adjacent<br>straight gyrus<br>(n = 48) |  |  |  |  |  |  |  |
|  | R | -0.078 | -0.017 | -0.167 | -0.102 | 0.015 | -0.066 |
|  | P | 0.598 | 0.908 | 0.257 | 0.492 | 0.918 | 0.654 |
| Superior<br>turbinate<br>(n = 48) |  |  |  |  |  |  |  |
|  | R | -0.384 | -0.270 | -0.126 | -0.120 | -0.352 | -0.281 |
|  | P | 0.007 | 0.064 | 0.394 | 0.418 | 0.014 | 0.053 |
| Middle turbinate<br>(n = 48) |  |  |  |  |  |  |  |
|  | R | -0.429 | -0.332 | -0.302 | -0.301 | -0.192 | -0.339 |
|  | P | 0.002 | 0.021 | 0.037 | 0.038 | 0.191 | 0.019 |
| Inferior<br>turbinate<br>(n = 48) |  |  |  |  |  |  |  |
|  | R | -0.394 | -0.303 | -0.318 | -0.264 | -0.194 | -0.257 |
|  | P | 0.006 | 0.036 | 0.028 | 0.070 | 0.187 | 0.078 |
| Optic nerve<br>(n = 53) |  |  |  |  |  |  |  |
|  | R | -0.054 | 0.156 | -0.172 | 0.107 | -0.174 | -0.047 |
|  | P | 0.702 | 0.265 | 0.219 | 0.447 | 0.212 | 0.736 |
| Corneoscleral<br>part (n = 53) |  |  |  |  |  |  |  |
|  | R | -0.024 | -0.057 | -0.027 | 0.020 | 0.086 | -0.124 |
|  | P | 0.862 | 0.686 | 0.848 | 0.884 | 0.540 | 0.377 |

T-MoCA: Telephone Montreal Cognitive Assessment.

### Relationship between CSF clearance and dyssomnia

The clearance function of superior turbinate was negatively correlated to the PSQI total scores, the clearance function of middle turbinate and inferior turbinate were negatively correlated to PSQI total scores, sleep quality and sleep latency, and the clearance function of corneoscleral part was negatively correlated to sleep habitual sleep efficiency (all *P* < 0.05; Table 2). No significant relationship was found between PSQI scores and the clearance function of parasagittal dura, adjacent straight gyrus and optic nerve (all *P* > 0.05; Table 2). The representative images in Fig. 5 show the relationship between the clearance function of superior turbinate, middle turbinate and inferior turbinate, and PSQI total scores.

**Table 2.** Correlations between sleep quality and percentage change of signal unit ratios from baseline to 39 hours in predefined seven locations.

|  |  | PSQI<br>total<br>scores | Sleep<br>quality | Sleep<br>latency | Sleep<br>duration | Habitual<br>sleep<br>efficiency | Sleep<br>disturbances | Use of<br>sleeping<br>medication | Daytime<br>dysfunction |
| --- | --- | --- | --- | --- | --- | --- | --- | --- | --- |
| Parasagittal dura (n = 53) |  |  |  |  |  |  |  |  |  |
| | $\rho$ | 0.097 | 0.089 | 0.138 | -0.001 | 0.132 | 0.049 | 0.098 | -0.052 |
|  | P | 0.488 | 0.528 | 0.325 | 0.996 | 0.348 | 0.726 | 0.483 | 0.711 |
| Adjacent straight gyrus (n=49) |  |  |  |  |  |  |  |  |  |
| | $\rho$ | 0.005 | 0.063 | -0.138 | -0.063 | -0.060 | -0.055 | 0.114 | 0.057 |
|  | P | 0.972 | 0.668 | 0.345 | 0.669 | 0.682 | 0.708 | 0.434 | 0.700 |
| Superior turbinate (n = 49) |  |  |  |  |  |  |  |  |  |
| | $\rho$ | 0.306 | 0.243 | 0.205 | 0.269 | 0.239 | 0.064 | -0.004 | 0.209 |
|  | P | 0.033 | 0.092 | 0.157 | 0.062 | 0.099 | 0.663 | 0.976 | 0.150 |
| Middle turbinate (n =49) |  |  |  |  |  |  |  |  |  |
| | $\rho$ | 0.375 | 0.314 | 0.400 | 0.145 | 0.187 | 0.057 | -0.044 | 0.237 |
|  | P | 0.008 | 0.028 | 0.004 | 0.319 | 0.199 | 0.697 | 0.764 | 0.101 |
| Inferior turbinate (n = 49) |  |  |  |  |  |  |  |  |  |
| | $\rho$ | 0.391 | 0.393 | 0.291 | 0.145 | 0.191 | -0.068 | 0.128 | 0.260 |
|  | P | 0.005 | 0.005 | 0.042 | 0.321 | 0.189 | 0.642 | 0.382 | 0.071 |
| Optic nerve (n = 55) |  |  |  |  |  |  |  |  |  |
| | $\rho$ | 0.110 | 0.163 | 0.113 | 0.151 | 0.227 | 0.020 | -0.134 | -0.042 |
|  | P | 0.424 | 0.233 | 0.411 | 0.271 | 0.095 | 0.887 | 0.329 | 0.763 |
| Corneoscleral part (n = 55) |  |  |  |  |  |  |  |  |  |
| | $\rho$ | 0.208 | 0.251 | 0.017 | 0.052 | 0.280 | 0.132 | -0.014 | 0.157 |
|  | P | 0.128 | 0.065 | 0.901 | 0.705 | 0.038 | 0.335 | 0.921 | 0.251 |

PSQI: Pittsburgh Sleep Quality Index

### Mediation of CSF clearance in the association of dyssomnia with cognitive decline

A total of 48 patients completed T-MoCA, PSQI and 2-dimensional T2-fluid attenuated inversion recovery at 39 hours. In the adjusted model, the PSQI total scores was negatively associated with the T-MoCA total scores (*β* = −0.235, *P* = 0.036; Table 3). The effect size of PSQI total scores on T-MoCA total scores after controlling for the clearance function of inferior turbinate was significantly reduced (change in *β* [bootstrap 95% confidence interval]: −0.082 [−0.220, −0.003]; Table 3). The mediator, clearance function of inferior turbinate, explained 35% of the association of PSQI total scores and T-MoCA total scores. The clearance function of superior and middle turbinate did not significantly mediate the association between PSQI total scores and T-MoCA total scores (change in *β* [bootstrap 95% confidence interval]: −0.047 [−0.149 to 0.020]; change in *β* [bootstrap 95%confidence interval]: −0.064 [−0.167 to 0.004]).

**Table 3.** Association of sleep quality with cognitive fuction mediated by the percentage change of signal unit ratios from baseline to 39 hours in inferior turbinate.

| Pathway | Unadjusted |  |  |  | Adjusted |  |  |  |
| --- | --- | --- | --- | --- | --- | --- | --- | --- |
| | $\beta$ | 95%CI | | P | $\beta$ | 95%CI | | P |
| a | 0.017 | 0.003 | 0.031 | 0.018 | 0.016 | 0.002 | 0.031 | 0.027 |
| b | -6.310 | -11.952 | -0.669 | 0.029 | -4.999 | -9.416 | -0.582 | 0.027 |
| c | -0.321 | -0.598 | -0.044 | 0.024 | -0.235 | -0.453 | -0.016 | 0.036 |
| c' | -0.213 | -0.496 | 0.070 | 0.136 | -0.153 | -0.374 | 0.068 | 0.170 |

a, the regression coefficient of the association of Sleep Quality Index (PSQI) total scores with the clearance function of inferior turbinate; b, the regression coefficient of association of the clearance function of inferior turbinate with Telephone Montreal Cognitive Assessment (T-MoCA) total scores, controlling for PSQI total scores; c, the regression coefficient of association of PSQI total scores with T-MoCA total scores; and c', the regression coefficient of association of PSQI total scores with T-MoCA total scores, controlling for the clearance function of inferior turbinate. Multivariable regression analysis for each pathway adjusted for the age and education years.

## Discussion

In the current study, we provided a method to simultaneously visualize the CSF clearance through three pathways: putative meningeal lymphatic pathway, peri-olfactory and peri-optic nerve pathways. The deep cervical lymph nodes were their potential CSF drainage way as the tracer appearance at the deep cervical lymph nodes were later than or at the same time point as above three pathways in 12 of 15 patients. The clearance function of putative peri-olfactory nerve pathway was impaired with aging, similar to putative meningeal lymphatic pathway. Importantly, CSF clearance function through the putative peri-olfactory inferior turbinate pathway explained 35% of the association of PSQI total scores and T-MoCA total scores, indicating its potential intervening mechanism involving dyssomnia and cognitive decline.

Rodents study have proved that the nasal region across the cribriform plate was the main CSF outflow site.^17^ Human postmortem also observed that Microfil dye injected into the CSF compartment located primarily in the subarachnoid space around the olfactory bulbs and cribriform plate.^18^ In all patients in our study, signal unit ratio in all three turbinates changed over time and in 86.3% patients, the tracer appeared at the adjacent straight gyrus before or at the same time point as nasal turbinates, indicating possible clearance of CSF to the turbinates via the cribriform plate along olfactory nerve. Although this finding is similar to previous results from PET study,^10^ significant change of signal intensity over time in the nasal turbinates was not observed in another study which also utilized intrathecal injection of tracer.^11^ The discrepancy may be due to the heterogeneity of disease, as half of the patients in their study were idiopathic normal intracranial pressure hydrocephalus and idiopathic intracranial pressure increased or decreased whose glymphatic function were impaired, potentially leading to limited drainage of CSF to the nasal turbinates, while our study mainly enrolled peripheral neuropathy patients with less disturbance in the CSF drainage. Additionally, the relatively limited sample size in their study reduced the statistical power, especially when the average signal unit ratio of turbinates changed slightly over time.

The three-layer sheath of the optic nerve continues with the meninges, and the dura mater and the arachnoid membrane fuse with the sclera at the distal end of the optic nerve.^19^ Lüdemann et al. observed that the arachnoid barrier layer was not continuous at the termination of the optic nerve and an outflow of tracer occurred in cat.^20^ Tracer was also found in the conjunctival lymphatics. Rodent study observed a dense plexus of tracer-filled lymphatic in the connective tissue inside the eyeball, and these lymphatics merged into a collecting vessel, tracking alongside the anterior facial vein to the mandibular lymph nodes.^21^ In humans, Jacobsen et al. found CSF enrichment within optic nerve by intrathecal injection of tracer, which also indicated the existence of glymphatic system in visual pathway.^22^ In current study, we directly observed the signal intensities both in middle intraorbital segment of the optic nerve and corneoscleral part changed over time, providing further evidence for the clearance of CSF through peri-optic nerve pathway. In addition, we found that in 88.7% patients, the tracer appeared at the optic nerve before or at the same time point as corneoscleral part, but there was no significant difference in the peak time point of the tracer in these two regions. We speculated that CSF might drain to the corneoscleral part via peri-optic space, but possible due to relatively long scan interval, it was not confirmed in our images. Further study with more observation time points and higher resolution MRI imaging is strongly needed to clarify this specific pathway.

In most patients (12/15), the tracer appeared at the deep cervical lymph nodes after or at the same time point as putative meningeal lymphatic pathway, putative peri-olfactory and peri-optic nerve pathways, supporting the potential drainage of CSF to deep cervical lymph node from cranium.^15,23^ The individual differences in the clearance might explain the unexpected results in the remaining 3 patients, as a certain pathway in some patients might contribute little to the overall CSF draining, which could not alter the peak time of deep cervical lymph node.

Multiple animal studies have demonstrated that CSF clearance function decrease with aging.^8,21,24^ Our previous study also found that the putative meningeal lymphatic pathway was impaired in aging human.^8^ In current study, we once again confirmed this conclusion with a larger sample size. Meanwhile, the similar relationship was also found in the putative peri-olfactory nerve pathway, which was accordant with the animal finding that less CSF drain to the nasal turbinates in aged mice.^17,25^ In our study, reduced clearance of the putative peri-optic nerve pathway with aging was not found. This indicates that the changes in the CSF clearance with aging may be inconsistent in the various pathways.

For the first time, we demonstrated that the dysfunction of CSF clearance from putative peri-olfactory nerve pathway was related to cognitive impairment. This finding is of importance as olfactory dysfunction is an important early manifestation in several neurodegenerative diseases such as Alzheimer’s disease.^26^ The pathological changes of Alzheimer’s disease widely emerge in the regions involved in olfactory information processing from the peripheral olfactory epithelium to the central entorhinal cortex prior to the symptomatic onset of cognitive impairment.^27,28^ Additionally, the odour identification performance is highly correlated with hippocampal volume in Alzheimer’s disease patients.^29^ Impaired peri-olfactory nerve pathway, associated with early olfactory dysfunction, may lead to the continuous deposition of abnormal protein and subsequent cognitive dysfunction. The predictive value of peri-olfactory nerve pathway on cognitive function may be worthwhile to test in different diseases.

The evidence for the relationship between CSF draining pathway and sleep in human is still limited, although studies have revealed that glymphatic influx function increased by 95% during sleep compared with wakefulness,^7^ and disrupted sleep architecture and depth may reduce the clearance of brain waste, as the efficacy of glymphatic function correlates with the prevalence of slow wave activity.^30^ Surprisingly, we found the impaired drainage of putative peri-olfactory nerve pathway, but unchanged drainage of parasagittal dura, in patients with chronic sleep disorders. Previously, unchanged tracer enrichment in parasagittal dura after total sleep deprivation of one night was also found,^31^ suggesting that molecular egress to parasagittal dura might be independent of acute or chronic sleep deprivation. Importantly, our further mediation analysis explained that impaired drainage of putative peri-olfactory nerve pathway might play an important role in the cognitive decline in patients with chronic sleep disorders, which also lends support to the connection between dyssomnia and cognitive impairment.^32^ Poor sleep quality increased amyloid *β* burden and promoted neuroinflammation, especially in hippocampal areas, resulting in cognitive impairment.^33,34^ Our findings thus provide evidence that peri-olfactory CSF clearance may be the important pathway for the drainage of abnormal proteins and inflammatory molecules in patient with poor sleep. This finding would have great clinical application for potential intervention of dementia by improving the peri-olfactory CSF clearance via the nasal drug delivery system that bypasses the blood-brain barrier and has good patient compliance.^35^

Our research has several limitations. First, the identification of putative meningeal lymphatic and peri-neural pathways were based on MRI, lacking corresponding pathological evidence. Second, we only obtained images at 3 time points after the intrathecal injection of the contrast agent with relatively long interval, which affected the accuracy of the peak time point. Third, the acquisition of patients’ cognitive data was not performed during hospitalization and was followed up by telephone after discharge. Fourth, the clearance function of the pathways could be affected by other factors, so the results on age, cognitive function and sleep variation should be interpreted with caution. Fifth, the clearance function of the pathways was simply regarded as the percentage change in signal unit ratio from baseline to 39 hours in regions. However, the clearance of the CSF tracer is a dynamic process which depends on the difference between influx and outflow speed of the tracer. Further studies with additional time points would help to understand the dynamic process of tracer clearance in more detail.

In conclusion, this study provides in vivo evidence of CSF clearance through putative meningeal lymphatic pathway, peri-olfactory and peri-optic nerve pathway in human and shows promise for dynamic MRI with intrathecal injection of contrast agent as a method to assess CSF clearance function through these pathways. This study also interprets the impaired peri-olfactory nerve clearance may explain the cognitive decline in patients with dyssomnia and indicates that it may be the important pathway for the drainage of abnormal proteins and inflammatory molecules in patient with poor sleep.

## Abbreviations

PSQI: Pittsburgh Sleep Quality Index
T-MoCA: Telephone Montreal Cognitive Assessment

