## Supplementary Table 1 for "Impaired peri-olfactory cerebrospinal fluid clearance mediated cognitive decline after dyssomnia"

**Supplemental Table 1. Basic clinical data**

|  | Total material |
| --- | --- |
| N | 85 |
| Age | 58 ± 14 |
| Female | 42 (49.4%) |
| Diabetes | 30 (35.3%) |
| Hypertension | 40 (47.1%) |
| Education years (n = 55) | 7.33 ± 4.10 |
| T-MoCA total scores (n = 54) | 14.17 ± 4.15 |
| Attention and calculation | 4.39 ± 1.63 |
| Language | 1.72 ± 1.09 |
| Abstraction | 0.63 ± 0.65 |
| Delayed recall | 2.09 ± 1.73 |
| Orientation | 5.33 ± 1.12 |
| PSQI total scores (n = 56) | 6.70 ± 4.20 |
| Sleep quality | 1.07 ± 0.89 |
| Sleep latency | 1.63 ± 1.15 |
| Sleep duration | 0.86 ± 0.94 |
| Habitual sleep efficiency | 0.63 ± 1.05 |
| Sleep disturbances | 0.98 ± 0.49 |
| Use of sleeping medication | 0.38 ± 1.00 |
| Daytime dysfunction | 1.14 ± 1.24 |

T-MoCA: Telephone Montreal Cognitive Assessment; PSQI: Pittsburgh Sleep Quality Index
