## Supplementary Table 2 for "Impaired peri-olfactory cerebrospinal fluid clearance mediated cognitive decline after dyssomnia"

**Supplemental Table 2. The peak time point of signal unit ratio at different locations among all patients**

| patient | Parasagittal dura | Optic nerve | Corneoscleral part | Adjacent straight gyrus | Superior turbinate | Middle turbinate | Inferior turbinate | Deep cervical lymph node |
| --- | --- | --- | --- | --- | --- | --- | --- | --- |
| #1 |  | 39h | 4.5h |  |  |  |  |  |
| #2 |  | 15h | 15h |  |  |  |  |  |
| #3 |  | 15h | 39h |  |  |  |  |  |
| #4 |  | 15h | 39h |  |  |  |  |  |
| #5 |  | 39h | 39h |  |  |  |  |  |
| #6 | 39h | 15h | 39h |  |  |  |  |  |
| #7 | 15h | 15h | 15h |  |  |  |  |  |
| #8 | 4.5h | 15h | 15h |  |  |  |  |  |
| #9 |  | 15h | 15h |  |  |  |  |  |
| #10 |  | 39h | 39h |  |  |  |  |  |
| #11 |  | 39h | 39h |  |  |  |  |  |
| #12 | 39h | 15h | 39h |  |  |  |  |  |
| #13 | 15h |  |  |  |  |  |  |  |
| #14 | 15h | 4.5h | 39h |  |  |  |  |  |
| #15 | 4.5h | 15h | 39h |  | 4.5h | 15h | 4.5h |  |
| #16 | 39h | 15h | 15h |  |  |  |  |  |
| #17 | 15h | 39h | 39h | 15h | 15h | 39h | 39h |  |
| #18 | 15h | 39h | 39h | 15h | 39h | 15h | 15h |  |
| #19 | 39h | 15h | 15h | 15h | 15h | 39h | 15h |  |
| #20 | 15h | 15h | 15h | 39h | 39h | 39h | 39h |  |
| #21 | 15h | 15h | 15h | 39h | 39h | 39h | 39h |  |
| #22 | 15h | 15h | 15h | 15h | 15h | 15h | 15h | 15h |
| #23 | 15h | 15h | 15h | 39h | 39h | 39h | 39h |  |
| #24 | 39h | 39h | 39h | 15h | 4.5h | 15h | 15h |  |
| #25 | 15h | 15h | 15h | 4.5h | 4.5h | 4.5h | 4.5h | 15h |
| #26 | 39h | 39h | 39h | 39h | 39h | 39h | 39h | 39h |
| #27 | 4.5h | 15h | 39h | 15h | 39h | 15h | 39h | 39h |
| #28 | 39h | 39h | 39h | 15h | 15h | 15h | 15h | 39h |
| #29 | 15h | 4.5h | 4.5h | 15h | 39h | 39h | 39h | 15h |
| #30 | 15h | 15h | 15h | 15h | 15h | 15h | 39h |  |
| #31 | 15h | 39h | 15h | 15h | 15h | 15h | 15h | 39h |
| #32 | 39h | 4.5h | 4.5h | 15h | 39h | 39h | 39h |  |
| #33 | 39h | 15h | 15h | 39h | 39h | 39h | 39h | 39h |
| #34 | 39h | 15h | 15h | 39h | 39h | 39h | 39h |  |
| #35 | 39h | 15h | 4.5h | 4.5h | 15h | 39h | 39h |  |
| #36 | 15h | 15h | 15h | 15h | 15h | 39h | 39h |  |
| #37 | 39h | 4.5h | 4.5h | 15h | 4.5h | 4.5h | 4.5h |  |
| #38 | 39h | 39h | 39h | 15h | 39h | 39h | 39h |  |
| #39 | 15h | 15h | 15h | 39h | 39h | 39h | 15h |  |
| #40 | 39h | 15h | 15h | 15h | 39h | 39h | 39h |  |
| #41 | 39h | 15h | 39h | 4.5h | 15h | 15h | 15h |  |
| #42 | 15h | 15h | 15h | 15h | 39h | 15h | 15h |  |
| #43 | 15h | 15h | 15h | 15h | 15h | 15h | 15h |  |
| #44 | 15h | 15h | 15h | 15h | 4.5h | 15h | 15h |  |
| #45 | 15h | 15h | 15h | 15h | 15h | 15h | 39h |  |
| #46 | 15h | 4.5h | 15h | 15h | 39h | 39h | 39h |  |
| #47 | 15h | 15h | 15h | 15h | 39h | 39h | 39h |  |
| #48 | 39h | 39h | 39h | 39h | 15h | 39h | 39h |  |
| #49 |  | 39h | 39h |  | 39h | 39h | 39h |  |
| #50 | 15h | 15h | 15h | 15h | 15h | 15h | 15h |  |
| #51 |  | 4.5h | 4.5h |  |  |  |  |  |
| #52 | 15h | 15h | 15h | 15h | 39h | 39h | 39h |  |
| #53 | 15h | 15h | 39h | 15h | 39h | 15h | 15h |  |
| #54 | 15h | 39h | 39h | 15h | 15h | 39h | 39h |  |
| #55 | 15h | 15h | 15h | 15h | 15h | 15h | 15h |  |
| #56 | 39h | 15h | 15h | 39h | 39h | 15h | 15h |  |
| #57 |  | 15h | 15h |  |  |  |  |  |
| #58 | 39h | 15h | 4.5h |  |  |  |  |  |
| #59 |  | 15h | 15h |  |  |  |  |  |
| #60 | 4.5h | 15h | 15h | 4.5h | 39h | 15h | 39h | 39h |
| #61 | 4.5h | 15h | 15h | 15h | 15h | 15h | 15h | 15h |
| #62 | 39h | 15h | 39h | 39h | 39h | 39h | 39h | 39h |
| #63 |  |  |  |  |  |  |  |  |
| #64 |  | 39h | 15h |  |  |  |  |  |
| #65 | 4.5h | 15h | 4.5h | 39h | 39h | 39h | 39h | 39h |
| #66 | 39h | 15h | 39h | 15h | 15h | 15h | 15h | 15h |
| #67 | 15h | 15h | 15h | 39h | 39h | 15h | 15h | 15h |
| #68 |  |  |  |  | 39h | 39h | 39h | 39h |
| #69 | 15h | 15h | 39h | 15h | 15h | 15h | 15h |  |
| #70 |  | 15h | 15h |  |  |  |  |  |
| #71 | 39h | 15h | 39h | 39h | 39h | 39h | 39h |  |
| #72 | 39h | 15h | 4.5h | 39h | 39h | 39h | 39h |  |
| #73 |  | 15h | 15h |  |  |  |  |  |
| #74 | 39h | 39h | 39h | 39h | 39h | 39h | 39h |  |
| #75 | 39h | 15h | 15h |  | 39h | 39h | 39h |  |
| #76 |  | 15h | 39h |  | 39h | 39h | 39h |  |
| #77 |  | 4.5h | 15h |  | 39h |  |  |  |
| #78 | 15h | 15h | 15h | 15h | 15h | 15h | 15h |  |
| #79 | 15h | 15h | 15h | 4.5h | 39h | 39h | 39h |  |
| #80 |  | 4.5h | 4.5h |  | 39h | 39h | 39h |  |
| #81 |  | 39h | 39h |  |  |  | 39h |  |
| #82 | 39h | 39h | 4.5h |  | 39h | 39h |  |  |
| #83 | 15h | 15h | 15h | 15h | 39h | 15h | 39h |  |
| #84 |  | 15h | 15h |  |  |  |  |  |
| #85 |  | 15h | 15h |  | 39h |  |  |  |

Missing value: the peak time point cannot be determined since the magnetic resonance sequence of the location is not scanned at a certain time point.
