## Supplementary Table 3 for "Impaired peri-olfactory cerebrospinal fluid clearance mediated cognitive decline after dyssomnia"

**Supplemental Table 3. Peak time point of cerebrospinal fluid tracer enrichment at different regions**

| Peak time point | Parasagittal  dura  (n = 62) | Adjacent  straight gyrus  (n = 51) | Superior turbinate (n = 60) | Middle turbinate (n = 58) | Inferior turbinate (n = 58) | Optic nerve  (n = 82) | Corneoscleral part  (n = 82) |
| --- | --- | --- | --- | --- | --- | --- | --- |
| 4.5h | 6 (9.7) | 5 (9.8) | 5 (8.3) | 2 (3.4) | 3 (5.2) | 8 (9.8) | 11 (13.4) |
| 15h | 31 (50.0) | 31 (60.8) | 19 (31.7) | 24 (41.4) | 20 (34.5) | 56 (68.3) | 43 (52.4) |
| 39h | 25 (40.3) | 15 (29.4) | 36 (60.0) | 32 (55.2) | 35 (60.3) | 18 (21.9) | 28 (34.2) |
| P | 0.354 | − | 0.006 | 0.004 | 0.002 | 0.450 | 0.990 |

The P-value refers to the paired samples Wilcoxon signed rank test of the other locations with the adjacent straight gyrus and the sample size was 51.
